# Sex-specific vulnerability to cannabinoid addiction-like behavior and associated miRNA signatures in a WIN 55,212-2 self-administration mouse model

**DOI:** 10.64898/2026.05.21.726674

**Authors:** Tatiana Gusinskaia, María Fernanda Ponce-Beti, Roberto Capellán, Edurne Gago-García, Noèlia Fernàndez-Castillo, Bru Cormand, Rafael Maldonado, Elena Martín-García

## Abstract

Cannabis is one of the most widely used psychoactive substances worldwide, and cannabis use disorder (CUD) is increasingly recognized as a major public health concern. Clinical evidence indicates that women may be particularly vulnerable to developing addiction, exhibiting a faster transition from initial drug use to loss of control and increased relapse vulnerability. However, females remain underrepresented in both preclinical and clinical research, limiting our understanding of the neurobiological mechanisms underlying this susceptibility. Here, we investigated sex differences in behavioral and epigenetic susceptibility to cannabinoid addiction using a mouse operant self-administration model with the synthetic cannabinoid agonist WIN 55,212-2. Female mice displayed increased addiction-like behavior, characterized by greater persistence of responding during drug-free periods and enhanced compulsive-like drug seeking compared to males. miRNA profiling in the medial prefrontal cortex (mPFC) identified a female-specific epigenetic signature associated with the addiction-like phenotype, including downregulation of mmu-miR-669j, mmu-miR-7036b-5p, mmu-miR-878-3p, and mmu-miR-7017-3p, together with upregulation of mmu-miR-3092-5p in addicted females. Functional enrichment analyses of predicted target genes revealed pathways related to synaptic organization, axon guidance, neurotransmission, and structural plasticity. Together, these findings demonstrate sex-dependent differences in vulnerability to cannabinoid addiction-like behavior and identify a specific miRNA signature in the mPFC associated with this phenotype, highlighting potential targets for the development of sex-specific therapeutic strategies.

## 1. INTRODUCTION

Drug addiction is a chronic relapsing disorder characterized by compulsive drug seeking and loss of control over drug intake despite adverse consequences (American Psychiatric Association, 2022). It represents a major global public health and socioeconomic burden (European Monitoring Centre for Drugs and Drug Addiction, 2022). Vulnerability to addiction is driven by complex interactions between genetic and environmental factors. Although psychiatric disorders are highly heritable and polygenic, phenotypic diversity arises from the convergence of genetic and environmental influences (Kenny, 2014). Epidemiological data reveal marked sex-dependent differences in substance use disorders. While men generally show higher rates of drug initiation and prevalence across most drug classes (UNODC, 2023), women exhibit a more rapid transition from initial exposure to loss of control, a phenomenon known as the telescoping effect, as well as higher relapse rates and more severe withdrawal symptoms (Towers et al., 2023). In the case of cannabis use disorder (CUD), the prevalence of which is increasing globally and influenced by expanding legalization (Hasin, 2018), differences are particularly pronounced, with women showing enhanced sensitivity to the reinforcing effects of cannabinoids and a faster progression toward severe addiction phenotypes (Khan et al., 2013; Towers et al., 2023).

Preclinical models are essential to dissect the neurobiological mechanisms underlying cannabinoid addiction. In rodents, the synthetic cannabinoid agonist WIN 55,212-2 provides a robust and reliable model of self-administration compared to Δ9-tetrahydrocannabinol (THC), which has a long pharmacokinetic half-life and often produces aversive effects and a narrow dose–response range (Maldonado, 2002; Valjent and Maldonado, 2000). WIN 55,212-2 acts as a short-acting and potent CB1 and CB2 receptor agonist and supports stable operant responding, enabling the study of the transition from controlled drug use to addiction-like behavior (Cajiao-Manrique et al., 2023b; Lefever et al., 2014; Martín-García et al., 2026).

The medial prefrontal cortex (mPFC) is a key brain region in this transition and plays a central role in executive control and the regulation of reward-seeking behavior. Dysregulation of the mPFC is a hallmark of addiction, contributing to impaired inhibitory control and reduced behavioral flexibility, thereby facilitating the shift toward compulsive drug use (Goldstein and Volkow, 2011; Koob and Volkow, 2016; Maldonado et al., 2021b). The mPFC exerts regulatory control over subcortical reward circuits, including the striatum, through tightly regulated glutamatergic signaling, which is critical for adaptive decision-making and behavioral flexibility (Koob and Volkow, 2016). At the molecular level, microRNAs (miRNAs) have emerged as critical epigenetic regulators of gene expression, capable of coordinating complex cellular responses to drugs of abuse and modulating entire gene networks, making them particularly relevant for remodeling synaptic organization, plasticity, and circuit-level adaptations associated with addiction vulnerability (Maldonado et al., 2021a).

Despite growing evidence supporting sex-dependent differences in cannabinoid-related behaviors, most preclinical studies have been conducted predominantly in males (Cajiao-Manrique et al., 2023a; Cajiao-Manrique et al., 2023b). Studies including both sexes suggest that females may exhibit enhanced sensitivity to cannabinoid reinforcement, including higher acquisition rates and increased drug-seeking behavior during abstinence (Fattore et al., 2007). However, the molecular mechanisms underlying these sex-dependent behavioral differences remain poorly understood.

To address this gap, we hypothesized that female vulnerability to cannabinoid addiction-like behavior is driven by alterations in prefrontal inhibitory control, leading to increased persistence of responding and compulsive-like drug seeking. Furthermore, we hypothesized that this behavioral phenotype would be associated with a sex-specific miRNA signature in the mPFC, reflecting dysregulation of gene networks involved in synaptic organization and neuronal plasticity. To test this, we combined a WIN 55,212-2 self-administration mouse model with the 3-criteria framework of addiction-like behavior and miRNA profiling in the mPFC.

## 2. METHODS

### 2.1. Animals

8-10-week-old C57BL/6J male (n=13) and female (n=18) mice (Charles River, France) were housed individually under controlled environmental conditions (21 ± 1 °C and 55 ± 10% humidity) with *ad libitum* access to water and a standard chow diet. All behavioral testing and self-administration sessions were performed during the dark phase of a reversed light cycle (lights off at 8.00 a.m. and on at 8.00 p.m.). Body weight was monitored weekly throughout the whole experimental period to ensure animal welfare. Animal procedures were conducted in strict accordance with the guidelines of the European Communities Council Directive 2010/63/EU and were approved by the local Ethical Committee for Animal Experimentation (Comitè Ètic d’Experimentació Animal-Parc de Recerca Biomèdica de Barcelona, CEEAPRBB, CEEA 1667-RML-22-0076-P1).

### 2.2. WIN 55, 212-2 Preparation

WIN 55, 212-2 ((R)-(+)-WIN 55,212-2 mesylate salt, Sigma-Aldrich, USA) was dissolved in 50 μL of Tween 80 (Sigma-Aldrich, USA) and subsequently diluted in sterile 0.9% saline. For the initial habituation, a priming dose of 0.1 mg/kg was administered via i.p. injection 24 h before the first operant session. For the intravenous self-administration sessions, the drug was prepared to deliver a dose of 12.5 μg/kg per infusion. All solutions were strictly protected from light and stored at room temperature throughout the duration of the experimental procedures.

### 2.3. Operant self-administration apparatus

Experiments were conducted in mouse operant conditioning chambers (model ENV-307A-CT, Med Associates Inc., Georgia, VT, USA) equipped with two nose-poke holes, with one randomly designated as the active hole and the other as the inactive hole. A house light was mounted on the chamber ceiling, and two stimulus lights were installed, one inside the active hole and the other above it. Nose-pokes in the active hole triggered the delivery of a single WIN 55,212-2 infusion according to the experimental schedule, accompanied by the activation of the stimulus lights inside and above the active hole. Nose-pokes in the inactive hole had no programmed consequences. Chambers were constructed from aluminum and acrylic and placed within sound- and light-attenuated boxes containing fans for ventilation and white noise. The chamber floor consisted of a stainless-steel bar grid capable of delivering electrical foot shocks during punishment tests. WIN 55,212-2 was administered in a volume of 23.5 µl over 2 seconds using a syringe mounted on a microinfusion pump (PHM-100A, Med-Associates, Georgia, VT, USA), connected via flexible polymer tubing (0.96 mm outer diameter, Portex Fine Bore Polythene Tubing, Portex Limited, Kent, UK) to a single-channel liquid swivel (375/25, Instech Laboratories, Plymouth Meeting, PA, USA) and subsequently to the mouse intravenous catheter.

### 2.4. Experimental design

Mice were anesthetized via intraperitoneal injection (0.1 ml per 10 g body weight) of ketamine hydrochloride (75 mg/kg, Ketamidor, Richter Pharma AG, Austria) and medetomidine hydrochloride (1 mg/kg, Domtor, Esteve, Spain) dissolved in 0.9% physiological saline. Indwelling intravenous catheters were subsequently implanted in the right jugular vein, as previously described (Mendizábal et al., 2006). After the surgery, animals were administered a solution for recovery and analgesia consisting of glucose serum (GlucosaVet, B. Braun Vet Care, Spain) and meloxicam (2 mg/kg, Metacam, Boehringer Ingelheim, Rhein), alongside a s.c. injection of the reversing agent atipamezole hydrochloride (2.5 mg/kg, Revertor, Virbac, Spain), all dissolved in sterile 0.9% physiological saline. Mice were housed individually and allowed to recover for 2 days with daily analgesic injections as described above.

After post-surgery recovery, the operant model was applied as recently described (Martín-García et al., 2026). Briefly, all animals received an ip injection of WIN 55, 212-2 24h before the start of self-administration daily sessions. The first five sessions followed a Fixed Ratio 1 (FR1) schedule, where each nose-poke resulted in one injection of WIN 55,212-2 at a dose of 12.5 μg/kg/infusion (Fig. 1A). The subsequent 5 sessions were changed to a Fixed Ratio 2 (FR2) schedule, requiring two active nose-pokes to obtain a single injection. Each session lasted 2 hours and 15 minutes and consisted of three periods: two 55-minute active periods separated by a 15-minute inactive drug-free period (DFP). During the DFP, responses resulted in no drug delivery, and the period was signaled by the house light being on. Sessions concluded after 50 reinforcers were delivered or after 125 min, whichever occurred first. If a mouse reached the 50-infusion limit, the threshold was increased to 100 reinforcers in the following session. After each session, catheters were flushed with 0.05 ml of 5% sodium heparin (Hospira, Pfizer) to maintain patency. Catheter patency was confirmed at the end of the self-administration sequence using a thiopental sodium test. Animals not exhibiting immediate signs of anesthesia were excluded. Mice that passed the test underwent 20 daily extinction sessions in the same chambers, during which WIN 55,212-2 and associated environmental cues were withheld.

**Figure 1.**
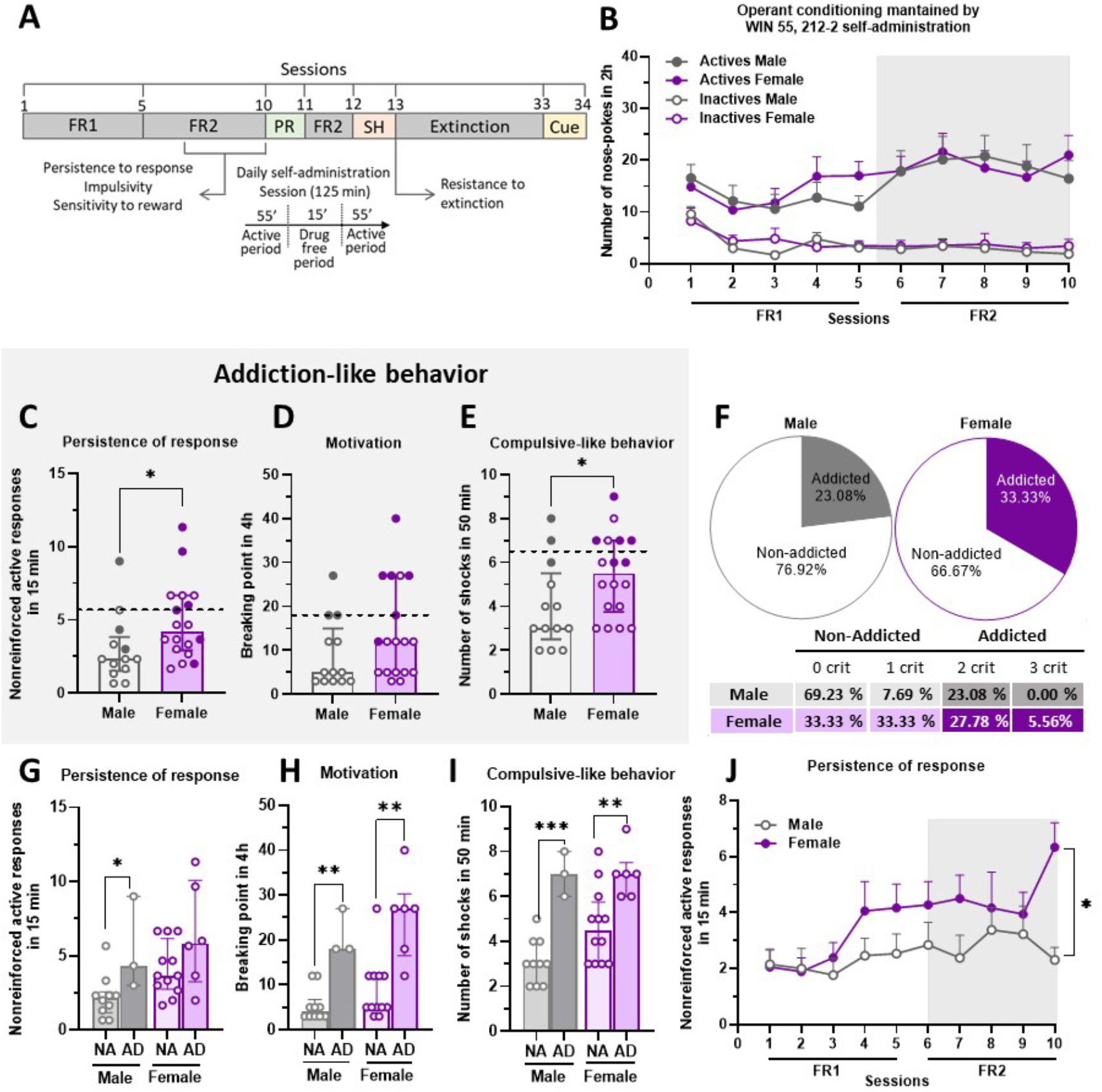
Female mice exhibit increased persistence of responding and compulsive-like behavior during WIN 55,212-2 self-administration. (A) Timeline of the experimental procedure, including five fixed ratio 1 (FR1) sessions, five fixed ratio 2 (FR2) sessions, a progressive ratio motivation test (PR), one FR2 re-establishment session, a compulsive-like behavior punishment test (SH), 20 extinction sessions, and a cue-induced reinstatement test (Cue). (B) Active and inactive nose-pokes performed by male and female mice during 2-h WIN 55,212-2 self-administration sessions under FR1 and FR2 schedules (mean ± SEM). (C–E) Evaluation of addiction-like behaviors in male and female mice (individual values with median and interquartile range (IQR)). (C) Persistence of responding (Mann–Whitney U test, *p < 0.05). (D) Motivation. (E) Compulsive-like behavior (Student’s t-test, *p < 0.05). The dashed horizontal line indicates the 75th percentile threshold used to classify animals as positive for a given addiction-like criterion. Addicted mice are represented by filled circles (gray for males and purple for females). (F) Percentage of mice classified as non-addicted (0– 1 criteria) or addicted (2–3 criteria) in male and female groups. (G–I) Behavioral performance of non-addicted (NA) and addicted (AD) mice separated by sex across the three addiction-like criteria (individual values with median and IQR). (G) Persistence of responding (Student’s t-test, *p < 0.05). (H) Motivation (Mann–Whitney U test, **p < 0.01). (I) Compulsive-like behavior (Student’s t-test, **p < 0.01, ***p < 0.001). Statistical significance is shown for NA vs AD comparisons within each sex. (J) Progression of persistence of responding during FR1 and FR2 WIN 55,212-2 self-administration sessions in male and female mice (mean ± SEM). Linear mixed model analysis revealed a significant session × sex interaction (*p < 0.05). Sample sizes were n = 13 for males and n = 18 for females (total n = 31).

Upon completion of the behavioral experiments, brains were rapidly extracted and coronal sections were obtained using a cryostat (Leica CM1950, Leica Biosystems Nussloch GmbH, Germany). The mPFC was microdissected and samples were immediately frozen and stored at -80°C for subsequent molecular analysis.

### 2.5. Addiction-like criteria

Following the FR1 and FR2 operant sessions of WIN 55,212-2, animals were evaluated across the three-core addiction-like behavioral hallmarks adapted from the DSM-5 substance use criteria (Cajiao-Manrique et al., 2023a; Cajiao-Manrique et al., 2023b; Martín-García et al., 2026).

#### 2.5.1. Persistence to response

Non-reinforced active nose-pokes were recorded during the 15-minute drug-free period (DFP). The values represent the mean score across the three consecutive FR2 sessions preceding the PR test.

#### 2.5.2. Motivation

The motivation test, using a progressive ratio (PR) schedule, was conducted in the self-administration chambers on the day following the fifth FR2 session. Over a four-hour period, the required number of nose-pokes to obtain a single injection increased exponentially according to the following sequence: 1, 2, 3, 5, 12, 21, 33, 51, 75, 90, 120, 155, 180, 225, 260, 300, 350, 410, 465, 540, 630, 730, 850, 1000, 1200, 1500, 1800, 2100, 2400, 2700, 3000, 3400, 3800, 4200, 4600, 5000, and 5500. The primary outcome measure was the breaking point, defined as the last completed series of nose-pokes required to obtain a reward. The session was terminated early if an animal remained inactive for more than one hour.

#### 2.5.3. Compulsive-like behavior

Compulsivity was assessed following an additional FR2 session to re-establish baseline behavior after the PR test. Mice were randomly assigned to a different self-administration chamber while maintaining their original active-hole side to avoid context-specific habituation. During a 50-minute session under an FR2 schedule, each nose-poke was paired with an aversive stimulus: an electric foot-shock (0.18 mA, 2-second duration). Following the first nose-poke, a 60-second countdown was initiated; the animal had to complete a second nose-poke within this window to obtain a WIN 55,212-2 infusion. If the second response was not completed within the time-out, the sequence was reset. The total number of shocks received during the session served as the quantitative measure of compulsivity.

### 2.6 Phenotypic traits considered as factors of vulnerability to addiction

To further characterize individual vulnerability factors, two additional phenotypic traits were assessed during the final stages of the acquisition phase.

#### 2.6.1 Impulsivity

Impulsivity was defined as the number of non-reinforced active nose-pokes performed during the 10-second time-out period following each infusion. Data were averaged over the three consecutive sessions preceding the PR test.

#### 2.6.2 Sensitivity to reward

The mean number of WIN 55,212-2 infusions earned across the three consecutive sessions prior to the PR test served as a quantitative measure of reward sensitivity.

### 2.7. Parameters related to craving

After 20 daily extinction sessions, two key parameters related to craving and relapse vulnerability were measured throughout the process.

#### 2.7.1. Resistance to extinction

The total number of non-reinforced active nose-pokes during the first 2-hour extinction session was used as a measure of resistance to extinction.

#### 2.7.2. Drug-seeking behavior

Following 20 extinction sessions, animals were exposed to a cue-induced reinstatement test (Cue). Two consecutive active nose-pokes triggered the cue light and the sound associated with drug delivery, although no infusion was administered. Drug-seeking was reported as the total number of active nose-pokes during the 2-hour session.

### 2.8. Animal classification

Mice were classified into “addicted” (AD) and “non-addicted” (NA) groups based on the number of addiction-like criteria met (motivation, persistence, and compulsivity). An animal was considered positive for a criterion if its score was at or above the 75th percentile of the total population distribution. Mice meeting 0 or 1 criterion were classified as non-addicted, while those meeting 2 or 3 criteria were classified as addicted.

### 2.9. miRNA extraction

The cerebral tissue was stored at -80ºC until extraction in 1.5 mL Eppendorf tubes. They were then disrupted and homogenized in a solution containing 10 μL of β-mercaptoethanol (SIGMA; Ref: M3148-100ML) per 1 mL of RLT Buffer from the AllPrep DNA/RNA/miRNA Universal Kit (Qiagen; 80224) using the Qiagen TissueRuptor and the corresponding disposable probes (Qiagen; Ref: 990890). 600 μL of the solution was used per sample. The samples were then centrifuged and transferred to the AllPrep DNA Mini spin column, which was placed in a 2 ml collection tube provided by the kit. They were centrifuged for 30 s at 20,000 x g. The flow-through was then transferred to a new 2 mL Eppendorf tube. 150 μl of chloroform (SIGMA; Ref: 1.02445) was then added to the sample, which was vortexed and centrifuged at 4°C for 3 min at 20,000 x g to separate the phases. The upper aqueous phase was then transferred to a 2 mL Eppendorf tube, and 80 μL of Proteinase K from the kit was added, followed by pipetting to mix. 350 μL of 100% ethanol (Mcquilkin; Ref: 85651.320) was then added and mixed. After a 10 min incubation at room temperature, 750 µl of 96–100% ethanol was added. The sample was then filtered through an RNeasy Mini spin column provided by the kit and centrifuged for 15 s at 20,000 x g; the flow-through was discarded. 500 µl of Buffer RPE (after adding 100% ethanol to reconstitute the concentrate) was added to the RNeasy Mini spin column, and the mixture was centrifuged for 15 s at 20,000 × g; the flow-through was discarded again. 10 µl DNase I stock solution (Qiagen; Ref: 79254) to 70 µl Buffer RDD. 80 µL of this mix is added per sample and incubated at room temperature for 15 min. 500 µL of the provided Buffer FRN is then added (after addition of isopropanol (SIGMA; Ref:109634) to reconstitute the concentrate). The samples were then centrifuged for 15 s at 20,000 x g. The flow-through was placed in the spin column again and centrifuged for the same duration at the same speed. The flow through was then discarded. 500 µL of the RPE Buffer was added to the RNeasy Mini spin column, and the column was centrifuged at the same speed for the same amount of time. The flow-through was discarded. 500 µl of 100% ethanol was added, and the tubes were centrifuged for 2 min at 20,000 x g. To eliminate any ethanol carryover, this centrifugation was repeated, adding the column to a new collection tube. The column was then placed in a 1.5 ml collection tube, and 50 µl of RNase-free water from the kit was added. The samples were then centrifuged for 1 min at 8000 x g to elute the RNA. Since a high RNA concentration was required, this last step was repeated using the eluate obtained. Samples were stored at -80º C.

### 2.10. miRNA sequencing and differential expression analysis

Libraries were prepared using the NEXTFLEX Small RNA-Seq Kit v4 for Illumina Platforms (ref. NOVA-5132, PerkinElmer) according to the manufacturer’s protocol. Briefly, the protocol comprised ligation of 3’ and 5’ adapters to the RNA molecules, reverse transcription to generate cDNA, and PCR amplification and indexing of cDNA molecules with adapters on both ends. PCR was performed using the UDI barcoded primer mix included in the kit, so final libraries had unique dual indexes. All purification steps were performed using the cleanup beads included in the kit. Final libraries were analyzed using Agilent Bioanalyzer HS DNA Assay (ref. 5067-4626) and quantified by qPCR using the KAPA Library Quantification Kit for Illumina (ROX Low) KK4873 (ref. 07960336001, Roche). The raw reads were preprocessed and aligned using an nf-core/smrnaseq (v.2.2.4, Docker profiler) (Ewels et al., 2020). Sequence alignment was performed using the GRCm38 assembly (GenBank accession: GCA_000001635.2, NCBI). Differential expression analysis was done by DESeq2 (Love et al., 2014). In order to discard those miRNAs with very low expression, we only considered the mature miRNAs showing at least ten reads across all the samples. To associate differentially expressed miRNAs with genes, the multiMiR package (Ru et al., 2014) was used, and only those predicted genes that were met in two or more databases were used for the following gene ontology (GO MF) and specific pathway (KEGG) analysis using the cluster Profiler package (Wu et al., 2021).

### 2.11. Statistical analysis

Statistical analysis of behavioral data was performed using IBM SPSS Statistics (v. 30.0). Normality was assessed using the Shapiro-Wilk test. Normally distributed data were analyzed using Student’s t-tests, while non-parametric data were analyzed using Mann-Whitney U tests. For longitudinal behavioral data, Linear Mixed Models (LMM) were employed. For transcriptomic data, differential expression was determined using the Wald test within DESeq2, and p-values were adjusted for multiple testing using the Benjamini-Hochberg False Discovery Rate (FDR) method. For GO and KEGG enrichment, Fisher’s Exact Test was applied to determine significant over-representation of functional categories.

## 3. RESULTS

### 3.1. Females exhibit increased vulnerability to addiction-like behaviors

To evaluate addiction-like behavior in male and female mice, animals underwent a WIN 55,212-2 self-administration protocol consisting of 34 separate sessions (Fig. 1A) (Martín-García et al., 2026). Both sexes successfully acquired an operant behavior characterized by a significant preference for the active nose-poke sensor over the inactive one across all sessions (Fig. 1B). No significant sex differences were observed in total drug intake.

Addiction-like behavior was assessed using three addiction-like criteria (Fig. 1C-E): Persistence of response (nose-pokes during drug-free period) (Fig. 1C), motivation (breaking point under a progressive ratio schedule) (Fig. 1D), and compulsive-like behavior (number of electric foot-shocks paired with the drug) (Fig. 1E). Female mice displayed significantly higher persistence (Mann-Whitney test U=58.00, Fig. 1C; p < 0.05) and compulsive-like responses despite adverse consequences (T-test t=2.091, Fig. 1E; p < 0.05) than males. Motivation under the progressive ratio schedule also tended to be higher in females, although differences did not reach statistical significance and the variance was broader (Fig. 1D). Specifically, when analyzing non-reinforced active nose-pokes during the drug-free period (persistence of response), female mice showed a progressive increase in seeking behavior that significantly diverged from males beginning at session 3 (Linear Mixed Model F(sex*session)=5.304, Fig. 1J; p < 0.05).

Animals were subsequently classified according to the number of addiction-like criteria fulfilled (Fig. 1F). Overall, 33.33% of females met the criteria for the addicted phenotype compared with 23.08% of males.

### 3.2. Behavioral profiling distinguishes addicted from non-addicted individuals

To evaluate the interaction between addiction classification and sex, behavioral patterns were compared between non-addicted (NA) and addicted (AD) individuals within each sex. In males, AD mice displayed greater persistence of responding compared with NA individuals (T-Test, t=2.592, Fig. 1G; p < 0.05). In contrast, female mice showed elevated persistence independently of addiction classification. Motivation and compulsivity differed significantly between AD and NA mice in both sexes. AD males and females showed increased motivation compared with NA animals (Mann-Whitney test U=0.01, Fig. 1H; p < 0.01 for male, Mann-Whitney test U=5.50, p < 0.01 for female). Similarly, compulsive-like behavior was significantly higher in AD mice of both sexes (T-test t=5.952, Fig. 1I; p < 0.001 for male, T-test t = 3.083, p < 0.01 for female). Interestingly, female NA mice also displayed higher persistence of responding (T-test t=-2.667, Fig. 1G, p < 0.05) and compulsive-like behavior (T-test t=-2.601, Fig 1I; p < 0,05) than NA males, suggesting that females generally exhibited higher responses for this criterion.

### 3.3. Addiction phenotype is associated with increased resistance to extinction

To assess the difficulty of extinguishing previous operant responses, mice underwent extinction training, in which active nose-pokes no longer resulted in the delivery of WIN 55,212-2. During 20 non-reinforced sessions (Fig. 2A), both sexes showed a gradual decrease in response. Nevertheless, female mice tended to show higher responding during the early stages of extinction (Fig. 2B). Specifically, AD mice of both sexes displayed significantly higher resistance to extinction on the first day compared to NA animals (T-test t=2.218, Fig. 2D, p < 0.05 for males, T-test t=3.135, p < 0.01 for females), suggesting that the addiction-like phenotype is strongly associated with an inability to inhibit drug-seeking behavior when the drug is unavailable.

**Figure 2.**
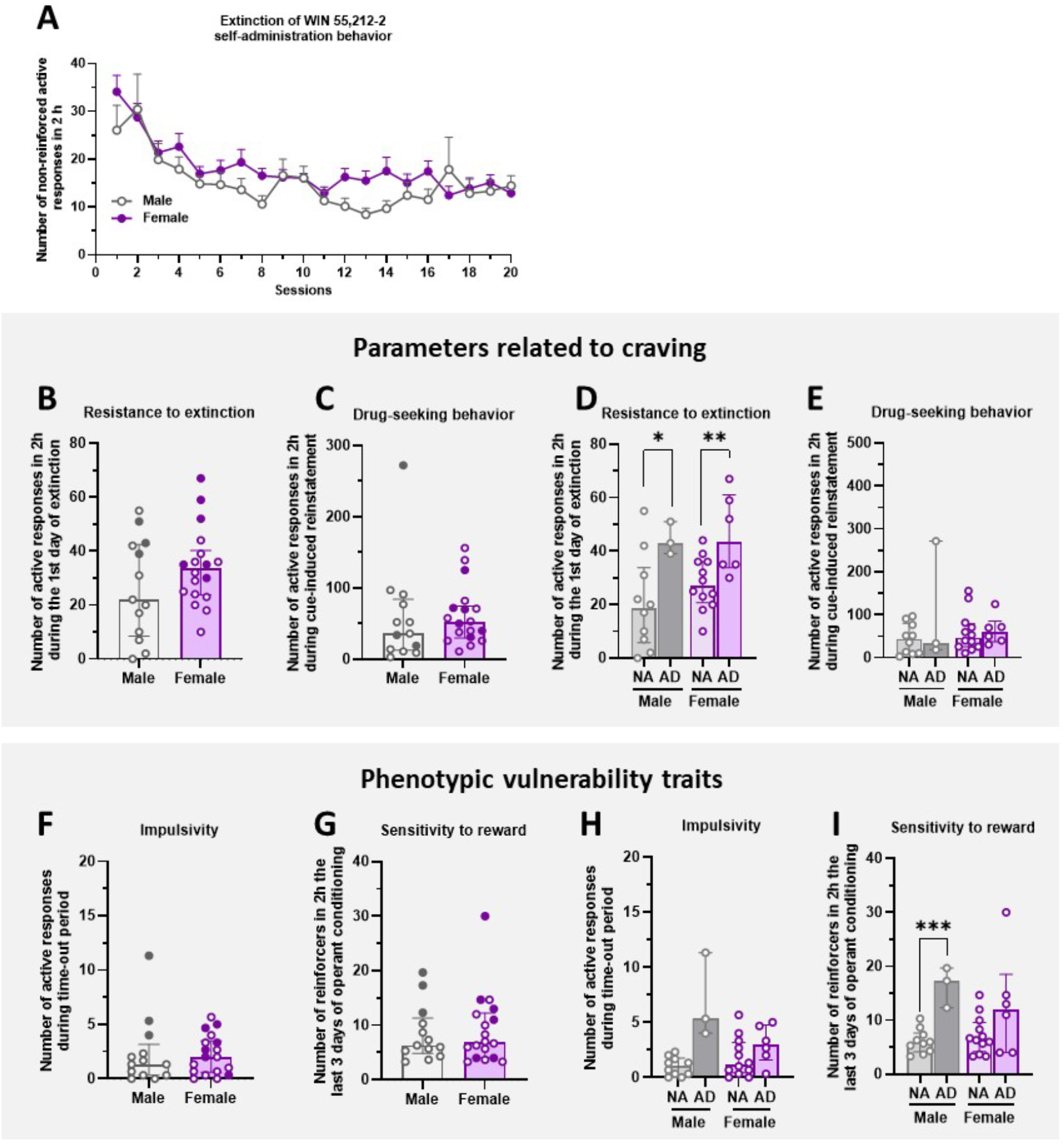
The addiction-like phenotype is associated with increased resistance to extinction and sex-dependent reward sensitivity. (A) Extinction profile of WIN 55,212-2 self-administration behavior in male and female mice during extinction sessions (mean ± SEM). (B,C) Behavioral measures related to craving (individual values with median and interquartile range (IQR)). (B) Resistance to extinction. (C) Drug-seeking behavior during cue-induced reinstatement. (D,E) Behavioral measures related to craving in non-addicted (NA) and addicted (AD) mice separated by sex (individual values with median and IQR). (D) Resistance to extinction (Student’s t-test, *p < 0.05 for males, **p < 0.01 for females). (E) Drug-seeking behavior. (F,G) Behavioral measures of phenotypic traits associated with vulnerability to addiction-like behavior (individual values with median and IQR). (F) Impulsivity. (G) Sensitivity to reward. (H,I) Behavioral measures of vulnerability-related traits in non-addicted (NA) and addicted (AD) mice separated by sex (individual values with median and IQR). (H) Impulsivity. (I) Sensitivity to reward (Student’s t-test, ***p < 0.001). Sample sizes were n = 13 for males and n = 18 for females (total n = 31).

Drug-seeking behavior was further evaluated in a cue-induced reinstatement test, as a measure of craving (Fig. 2C). Although no significant sex differences were detected in drug-seeking behavior, AD male mice attended to display a higher number of responses than NA males, albeit with substantial variability within the AD male group (Fig. 2E).

### 3.4. Phenotypic traits reveal sex-specific patterns in reward sensitivity

To identify phenotypic traits that can be aligned to vulnerability to drug addiction, impulsivity (Fig. 2F), and sensitivity to reward (Fig. 2G) were measured. No significant sex differences were observed for either measure. As with drug-seeking behavior, AD male mice showed a trend toward increased impulsivity compared with NA males, although variability within the AD group was high (Fig. 2H). In contrast, AD male mice displayed significantly increased response in the last three days of self-administration sessions relative to NA males (T-test t=6.158, Fig. 2I, p<0.001), while females exhibited more homogeneous levels of responding across addiction classifications, which can be considered as a sign of “telescopic” effect in the female group.

### 3.5. Principal component analysis reveals a behavioral dimension associated with addiction-like phenotype

To evaluate whether behavioral variables segregated according to sex or addiction-like classification, we performed a principal component analysis (PCA). The first two principal components explained 66% of the total variance (PC1: 51%; PC2: 15%). Animals were separated primarily by addiction-like phenotype along the PC1 axis rather than by biological sex (Fig. 3A). Specifically, AD mice clustered toward positive PC1 values, whereas NA animals clustered toward negative values.

**Figure 3.**
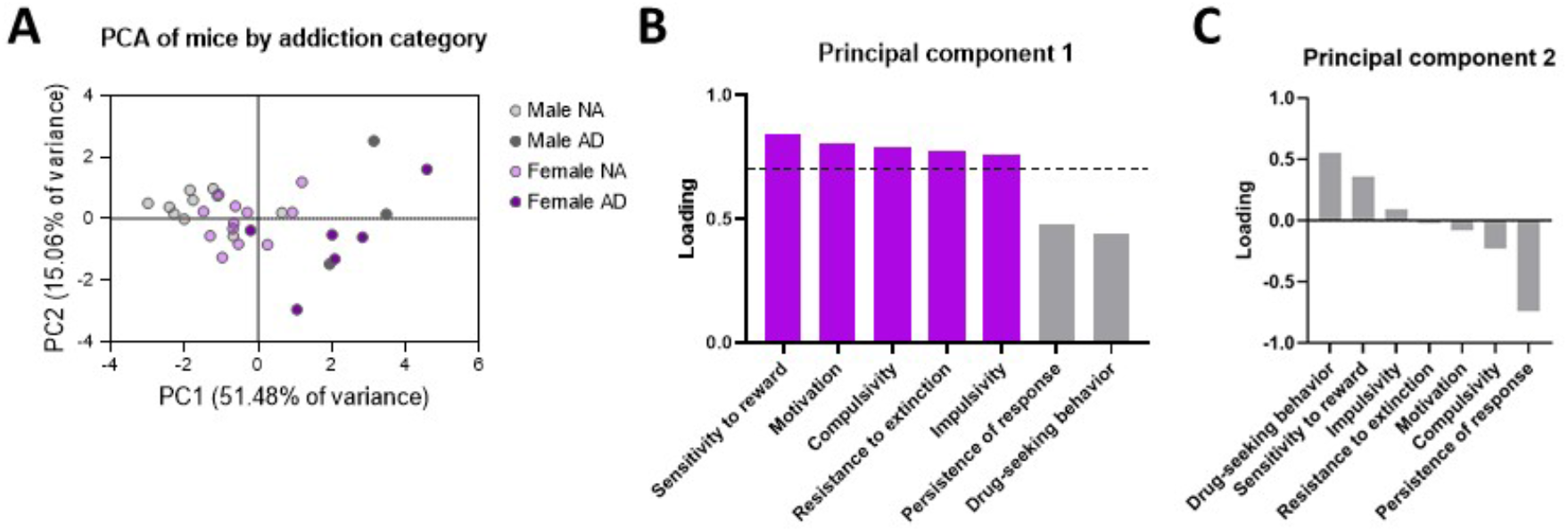
Principal component analysis reveals distinct behavioral dimensions associated with addiction-like phenotype. (A) Distribution of individual mice according to the first two principal components (PC1 and PC2), which together accounted for 66.54% of the total variance (PC1: 51.48%; PC2: 15.06%). Animals are color-coded according to sex (male, female) and addiction-like classification (addicted, AD; non-addicted, NA). Separation along PC1 primarily reflected addiction-like phenotype rather than biological sex, with addicted mice shifted toward positive PC1 values. (B) Variable loadings contributing to PC1. Sensitivity to reward, motivation, compulsive-like behavior, resistance to extinction, and impulsivity exhibited the strongest positive loadings (> 0.7 threshold, dashed line), indicating that PC1 captures a major addiction-related behavioral dimension. (C) Variable loadings contributing to PC2. Persistence of responding exhibited a strong negative loading, whereas drug-seeking behavior and sensitivity to reward showed weaker contributions below the 0.7 threshold, suggesting that PC2 reflects a partially dissociable behavioral dimension associated with persistence-related traits.

Analysis of variable loadings (Fig. 3B) showed that motivation, compulsive-like behavior, sensitivity to reward, and resistance to extinction contributed most strongly to PC1, indicating that this component represents a major addiction-related behavioral dimension. In contrast, PC2 was characterized by a strong negative loading for persistence of responding and weaker positive contributions from drug-seeking behavior, suggesting that these traits may represent partially dissociable behavioral dimensions.

### 3.6. Transcriptomic profiling reveals a female-specific miRNA signature of addiction

To investigate molecular alterations associated with addiction development, miRNA sequencing was performed in the mPFC. Differential expression analysis identified a distinct miRNA profile in AD female mice compared with the remaining experimental groups (p adjusted < 0.01, |log2FoldChange| > 1, Fig. 4A). Specifically, four miRNAs were significantly downregulated (*mmu-miR-669j, mmu-miR-7036b-5p, mmu-miR-878-3p, mmu-miR-7017-3p*) and one upregulated (*mmu-miR-3092-5p*) in AD females. In contrast, other commonly expressed miRNAs, including members of the let-7 and miR-1 families, remained stable (NS - non-significant), suggesting that the observed changes represent a selective molecular signature associated with female vulnerability to cannabis addiction in females.

**Figure 4.**
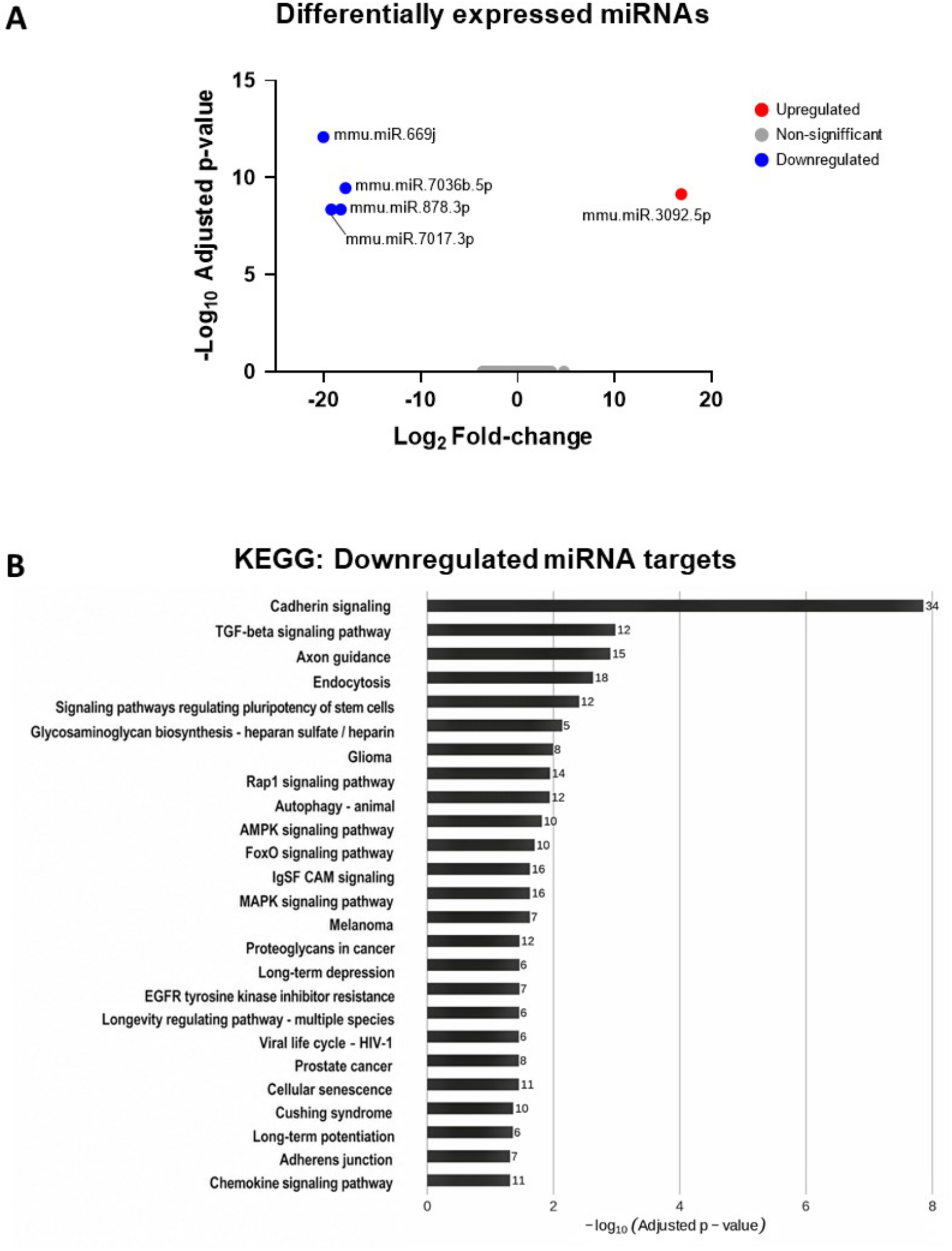
Differential miRNA expression and functional enrichment analyses in addicted female mice. (A) Volcano plot showing differentially expressed miRNAs in addicted female mice compared with the remaining experimental groups. Significantly upregulated and downregulated miRNAs are highlighted in red and blue, respectively, whereas non-significant miRNAs are shown in gray (thresholds: |log2 fold-change| > 1 and adjusted p-value < 0.05). (B) Top 25 enriched KEGG pathways associated with predicted target genes of downregulated miRNAs. For panel B, pathways and biological processes were ranked according to adjusted p-values. The x-axis represents statistical significance (−log10 adjusted p-value), and the numbers shown beside each bar indicate the number of target genes associated with each pathway or GO term.

### 3.7. Dysregulated miRNAs target structural plasticity and memory pathways

To elucidate the molecular functions potentially associated with the dysregulated miRNAs, we performed Gene Ontology (GO) and KEGG pathway enrichment analyses for genes targeted by both downregulated (*mmu-miR-669j, mmu-miR-7036b-5p, mmu-miR-878-3p, mmu-miR-7017-3p*) and upregulated miRNAs (*mmu-miR-3092-5p*). KEGG pathway analysis identified enrichment in pathways associated with TGF-beta signaling, axon guidance, and MAPK signaling (Fig. 4B). Although KEGG enrichment did not reach statistical significance, several predicted targets were associated with pathways related to addiction: “amphetamine addiction” and “dopaminergic synapse” (p adj = 0.09). Predicted targets of downregulated miRNAs were enriched for processes related to neuronal architecture and synaptic signaling. Top GO terms included “positive regulation of cell projection organization”, “regulation of neurogenesis”, and “regulation of synapse structure or activity”, as well as pathways related to “axon guidance”, “neuron projection guidance”, and “regulation of neurotransmitter secretion” (Fig. 5A). Predicted targets of the upregulated miRNA, *mmu-miR-3092-5p*, were enriched in processes related to synapse organization, synaptic activity, and cognitive functions such as learning and memory (Fig. 5B).

**Figure 5.**
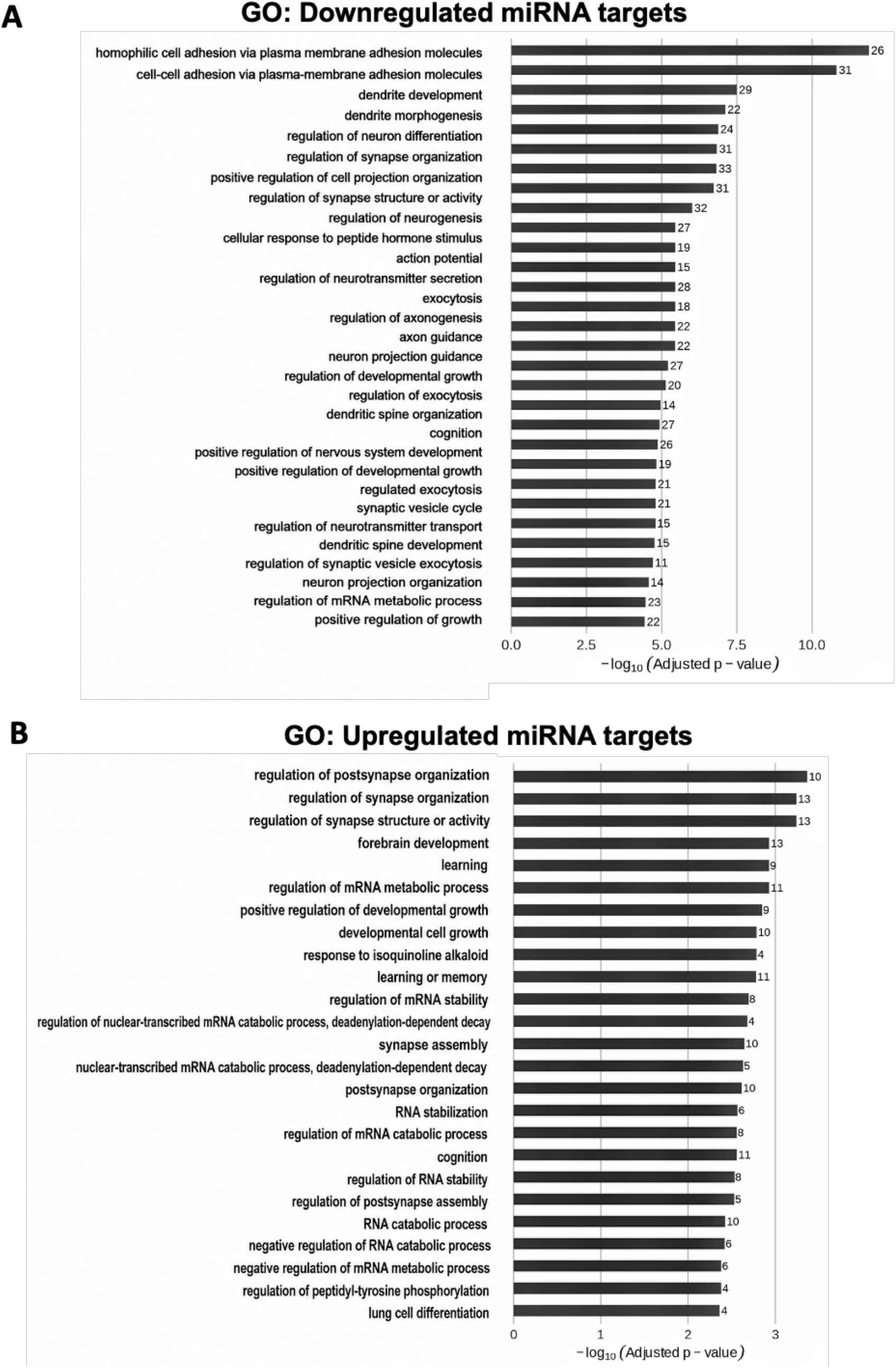
**Top enriched Gene Ontology (GO) molecular functions** associated with predicted target genes of (A) downregulated and (B) upregulated miRNAs. For panels A-B, pathways and biological processes were ranked according to adjusted p-values. The x-axis represents statistical significance (−log10 adjusted p-value), and the numbers shown beside each bar indicate the number of target genes associated with each molecular function or GO term.

## 4. DISCUSSION

The present findings suggest that female vulnerability to cannabinoid addiction-like behavior is associated with miRNA-mediated alterations in pathways involved in synaptic organization, neuronal plasticity, and reward-related circuitry, potentially contributing to impaired inhibitory control and compulsive-like drug seeking.

Cannabis use disorder (CUD) represents a significant global health challenge with increasing prevalence as legalization expands across different regions (UNODC, 2023). Epidemiological data consistently highlight pronounced sex-dependent differences in its progression. Although men generally exhibit higher rates of cannabis initiation, clinical studies indicate that women may progress more rapidly toward dependence and severe addiction phenotypes. In the present study, we investigated the neurobiological basis of these sex differences using an intravenous WIN 55,212-2 self-administration model in C57BL/6J mice, the 3-criteria framework of addiction-like behavior, and miRNA profiling in the mPFC. This area was selected for analysis due to its critical role in executive control and the top-down regulation of reward-seeking behavior, as these functions are often compromised in individuals with addiction (Cajiao-Manrique et al., 2023a; Domingo-Rodriguez et al., 2020; García-Blanco et al., 2022).

Our results demonstrate that female mice exhibit increased vulnerability to specific dimensions of cannabinoid addiction-like behavior, particularly enhanced persistence of responding during drug-free periods and greater compulsive-like drug seeking despite punishment. Thus, the proportion of females meeting the criteria for an addiction-like phenotype was 10% higher than that of males, supporting the existence of sex-dependent susceptibility to addiction. These behavioral alterations were associated with a distinct miRNA signature in the mPFC, suggesting that sex-specific molecular adaptations in prefrontal circuits may contribute to the observed vulnerability. Importantly, the behavioral profile observed in females was not explained by higher drug intake or by a general increase in motivation for the drug. While motivation measured under a progressive ratio schedule did not differ significantly between sexes, female mice displayed significantly greater persistence of responding during drug-free periods and greater resistance to punishment than male mice, indicating enhanced compulsive-like drug-seeking (Cajiao-Manrique et al., 2023b; Fattore et al., 2007; Maldonado et al., 2021b). These findings suggest that female vulnerability to cannabinoid addiction-like behavior is primarily associated with impaired inhibitory control and persistence of drug seeking despite negative consequences, rather than increased cannabinoid reinforcement or motivation. This interpretation is further supported by the principal component analysis, which revealed that animals segregated primarily according to addiction-like phenotype rather than sex. Notably, female mice displayed a behavioral distribution shifted toward the addiction-like cluster compared with males, consistent with an increased vulnerability profile. Compulsive-like behavior contributed strongly to PC1, the principal addiction-related behavioral dimension of loss of control, whereas persistence of responding was more strongly associated with PC2, suggesting that these criteria may represent partially dissociable dimensions of addiction-like behavior. Importantly, females exhibited significantly increased compulsivity and persistence of responding compared with males, indicating that female vulnerability extends across multiple behavioral dimensions linked to impaired inhibitory control.

To investigate the molecular correlates of this vulnerability, we analyzed miRNA expression in the mPFC and identified a distinct epigenetic female signature associated with the addicted phenotype. Addicted females exhibited downregulation of mmu-miR-669j, mmu-miR-7036b-5p, and mmu-miR-878-3p, together with upregulation of mmu-miR-3092-5p. Given the central role of the mPFC in executive control, these findings suggest that sex-specific epigenetic regulation may contribute to the impaired inhibitory control and compulsive-like behavior observed in females. Importantly, these miRNA alterations provide a potential mechanistic link between prefrontal dysfunction and the behavioral vulnerability observed in females.

The significant downregulation of *miR-669j* and *miR-878-3p* in addicted females is particularly relevant when considering the enrichment of pathways involved in the regulation of post-synapse organization. Since miRNAs typically act as negative regulators of gene expression, their decrease likely leads to the overexpression of target genes, such as *Erbb4* and *Kalrn. Erbb4* is a critical regulator of GABAergic interneuron function, and its dysregulation in the mPFC has been shown to impair top-down regulation and prefrontal inhibitory control (Tan et al., 2018). We hypothesize that this molecular shift may facilitate the persistence of responding and compulsive-like behavior observed in females, as altered GABAergic signaling in the mPFC may disrupt the excitatory-inhibitory balance and impair the capacity of prefrontal circuits to exert top-down control over subcortical reward regions, such as the ventral and dorsal striatum. This dysfunction would compromise the ability to suppress drug-seeking behavior when the drug is unavailable or when adverse consequences are present. Similarly, the dysregulation of *Kalrn*, which is essential for dendritic spine plasticity, suggests a possible pathological remodeling of prefrontal circuits that may weaken prefrontal– striatal connectivity and may contribute to the transition toward compulsive-like behavior (Schaefer et al., 2010).

On the other hand, upregulation of mmu-miR-3092-5p is predicted to reduce the expression of genes involved in homophilic cell adhesion, dendritic development, and cognitive processes, such as learning and memory. Functional enrichment analysis revealed that the targets of this miRNA are primarily associated with synapse organization and activity, processes essential for the encoding and updating of reward-related memories. In the context of substance use disorders, addiction is increasingly conceptualized as a disorder of maladaptive learning (Everitt and Robbins, 2016; Koob and Volkow, 2016). Therefore, miR-3092-5p-mediated repression of these genes may impair the molecular mechanisms underlying cognitive flexibility and adaptive memory updating (Goldstein and Volkow, 2011). Among these targets, the possible downregulation of *Cadm2*, a predicted target of mmu-miR-3092-5p, could be of particular interest, as genetic variants in this gene have been strongly associated with risky behaviors and CUD in human GWAS studies (Arends et al., 2021). Also, genetic variants in *CADM2* were found to exert pleiotropic effects between several substance use disorders (including cannabis, tobacco, opioids and alcohol) and attention-deficit/hyperactivity disorder (Koller et al., 2024). Additionally, members of the protocadherin family, such as *Pcdh10* and *Pcdh19*, further support the possible involvement of cell-adhesion mechanisms (Missler et al., 2012; Südhof, 2018). As cell adhesion molecules are critical for maintaining synaptic stability, their reduction may lead to a state of synaptic destabilization in the female mPFC that may disrupt the structural plasticity required for adaptive behavioral regulation (Kalivas, 2009; Lüscher and Malenka, 2011). This synaptic destabilization may further weaken prefrontal control over subcortical reward circuits, contributing to the persistence of maladaptive drug-seeking behavior.

This possible structural instability may be further compounded by the dysregulation of genes involved in glutamatergic homeostasis, as suggested by our target prediction and enrichment analyses. Among these, SLC1A2, which encodes the glutamate transporter GLT-1 (EAAT2), represents a potential candidate. SLC1A2 is a major regulator of glutamate homeostasis in the neocortex, a process essential for maintaining the synaptic precision required for adaptive learning and memory formation (Bjørnsen et al., 2014). In addictive states, reduced SLC1A2-mediated glutamate clearance has been associated with increased extracellular glutamate levels, which may disrupt the balance between tonic and phasic glutamatergic signaling. This imbalance can reduce the signal-to-noise ratio of prefrontal glutamate transmission, ultimately impairing the ability of mPFC circuits to exert precise top-down control over striatal reward pathways (Kalivas, 2009; Knackstedt et al., 2010). In this context, glutamate dysregulation may contribute to reduced cognitive control and behavioral flexibility, rather than a simple increase in excitatory drive. The clinical association of SLC1A2 variants with elevated cortical glutamate and increased psychiatric vulnerability (Veldic et al., 2019; Yahya et al., 2023) further supports the idea that SLC1A2 dysregulation may be a downstream consequence of the miRNA alterations observed here, potentially contributing to impaired inhibitory control in females.

Although KEGG analysis for mmu-miR-3092-5p did not reach statistical significance, its predicted targets play an important role in pathways related to the dopaminergic synapse and amphetamine addiction. In this context, mmu-miR-3092-5p-mediated downregulation of genes involved in synaptic organization and memory processes may contribute to a weakening of prefrontal circuit function, reducing the capacity to exert top-down control over subcortical reward systems. Such alterations may underlie the increased resistance to punishment and persistence of drug-seeking behavior observed in females. Furthermore, the identification of factors such as *Ago2* and *Eif4e2* among the predicted targets of mmu-miR-3092-5p suggests that this miRNA may contribute to broader alterations in post-transcriptional regulation associated with synaptic plasticity (García-Blanco et al., 2022; Schaefer et al., 2010).

In conclusion, our findings indicate that sex is an important factor influencing vulnerability to cannabinoid addiction-like behavior, with female mice exhibiting a greater propensity for persistence and compulsive-like drug seeking. This behavioral phenotype does not appear to be driven by enhanced motivation for the drug, but rather by deficits in inhibitory control mechanisms that may reflect prefrontal dysfunction. This increased vulnerability is associated with a distinct miRNA-mediated transcriptomic signature in the mPFC, characterized by downregulation of miRNAs involved in synaptic organization and upregulation of miRNAs targeting genes involved in synaptic stability, learning, and memory. Together, these alterations point to a state of prefrontal circuit dysregulation that may promote the persistence of maladaptive drug-seeking behaviors. Importantly, these findings highlight the mPFC as a critical hub underlying sex-specific vulnerability to cannabinoid addiction and provide a mechanistic framework for the development of targeted therapeutic strategies.

## Acknowledgements

This work was supported by Spanish “Ministerio de Ciencia, Innovación y Universidades, Agencia Estatal de Investigación (AEI)” (PID2020-120029GB-I00/MICIN/AEI/10.13039/501100011033, RD21/0009/0019 and PDI2023-1511680B-C21), the Spanish “Instituto de Salud Carlos III, RETICS-RTA” (#RD12/0028/0023), the “Generalitat de Catalunya, AGAUR” (#2020 SGR), “ICREA-Acadèmia” (#2025) and the Spanish “Ministerio de Sanidad, Servicios Sociales e Igualdad”, “Plan Nacional Sobre Drogas of the Spanish Ministry of Health” (#PNSD-2022) to R.M., “la Caixa Health” LCR/PR/HR22/5240017 to R.M. and E.M-G., “Plan Nacional Sobre Drogas of the Spanish Ministry of Health” (#PNSD-2019I006, #PNSD-2023I040) and Spanish “Ministerio de Ciencia e Innovación” (ERA-NET) PCI2021-122073-2A to E.M-G. Funding supporting this study was provided by the Spanish ‘Ministerio de Ciencia, Innovación y Universidades’ (PID2024-158634OB-I00, to NFC and BC), ‘Generalitat de Catalunya/AGAUR’, (2021-SGR-01093, to NFC and BC), ICREA Academia 2021 (to BC), ‘Fundació La Marató de TV3 ′ (202218-31, to BC) and ‘Ministerio de Sanidad, Servicios Sociales e Igualdad/Plan Nacional Sobre Drogas’ (PNSD-2020I042 and PNSD-2024I056, to NFC).

We are very grateful to R. Martín and F. Porrón for their technical support. Figure 1. A was created with BioRender.

## Author contributions

E.M.-G. and R.M. conceived and designed the behavioral studies. T.G., M.F.P.-B., and R.C. performed the behavioral experiments and statistical analyses under the supervision of E.M.-G. and R.M. The intravenous catheterization surgeries were performed by E.M.-G. with assistance from T.G., M.F.P.-B., and R.C. The RNA extraction was performed by E.G.-G. under the supervision of N.F.-C. and B.C. Bioinformatic analyses were performed by T.G. with input from E.M.-G., N.F.-C., B.C., and R.M. Next, T.G., M.F.P.-B., R.C., E.M.-G., and R.M. wrote the manuscript. E.G.-G., N.F.-C., and B.C. contributed to manuscript editing and revision. Finally, R.M. and E.M.-G. critically revised the manuscript with input from all authors.

## Data availability

Individual data points are graphed in the main figures. All the relevant data that support this study are available from the corresponding author to any interested researcher upon reasonable request.

## Conflict of interest

The authors declare no conflict of interest.

## Notes

### Competing Interest Statement

The authors have declared no competing interest.

